# Ubiquitin-specific peptidase 20 constrains endothelial cell activation and angiogenic sprouting through NF-κB inhibition

**DOI:** 10.1101/2025.05.20.655129

**Authors:** Bipradas Roy, Jiao-hui Wu, Richard Jiang, Annie Bao, Neil J. Freedman, Sudha K. Shenoy

## Abstract

Nuclear factor-κB (NF-κB) mediates inflammation-driven angiogenesis, which promotes the growth of atherosclerotic plaques and tumors. The deubiquitinase ubiquitin-specific peptidase 20 (USP20) suppresses NF-κB activation in vascular smooth muscle cells (SMCs) and attenuates atherosclerosis. Phosphorylation of USP20 at Ser334 increases NF-κB signaling in SMCs. However, the role of USP20 in endothelial cells (ECs) remains undefined. We therefore tested whether USP20 activity diminishes NF-κB signaling in ECs and thereby diminishes angiogenesis. Cytokine-induced NF-κB activity was elevated in primary ECs isolated from *Usp20*^-/-^ mice as compared with ECs from wild-type (WT) mice. Concordantly, cytokine-induced NF-κB activity was elevated in mouse coronary ECs (MCECs) expressing dominant-negative USP20 (USP20-DN) or phospho-mimetic USP20 (USP20-S334D), but *blunted* in MCECs expressing WT USP20 (USP20-WT) or phospho-resistant USP20 (USP20-S334A). MCEC migration and spheroid sprouting/angiogenesis were increased with overexpression of USP20-DN or USP20-S334D, but decreased with overexpression of USP20-WT or USP20-S334A. Angiogenesis assessed by the aortic ring assay was significantly increased in *Usp20*^-/-^ aortas and suppressed by TPCA-1, an inhibitor of NF-κB signaling. Angiogenesis was augmented in *Usp20*-S334D aortic rings but reduced in *Usp20*-S334A aortic rings. By screening known angiogenic factors, we identified matrix metalloproteinase 3 (MMP-3), a transcriptional target of NF-κB, as a gene that is also regulated by USP20 expression. Inhibiting MMP-3 reduced angiogenic sprouting in the *Usp20*^-/-^ mouse aortic rings. We conclude that USP20 expression inversely correlates with the extent of angiogenesis, and that inhibiting USP20 Ser334 phosphorylation could be a useful strategy to constrain inflammation-driven angiogenesis in pathological conditions like solid tumor malignancies and atherosclerosis.

## INTRODUCTION

In endothelial cells (ECs), activation of the proinflammatory transcription factor NF-κB plays an integral role in the formation and maturation of new blood vessels (1,2)—processes critical not only to normal development and wound healing but also to the progression of solid tumor malignancies (3–6) and atherosclerosis (7,8), among other pathologies. The activation of NF-κB signaling depends on IκB kinase-β activation, which is regulated by a cascade of enzymatic steps that activate proteins through lysine-63-linked, reversible polyubiquitination (9,10). Proinflammatory cytokine-mediated activation of tumor necrosis factor receptor (TNFR), Toll-like receptor-4 (TLR4) or interleukin-1 receptor (IL-1R) triggers lysine-63-linked ubiquitination of the downstream adaptor protein TNFR-associated factor (TRAF) (10). As part of its activation process, TRAF autoubiquitinates and subsequently ubiquitinates NF-κB essential modulator (NEMO) and the TAB2 subunit of the TAK1 kinase complex. Ubiquitinated TAK1 phosphorylates and activates the inhibitor of κB kinase-β (IKKβ), which in turn phosphorylates not only the p65 subunit of NF-κB but also the inhibitor of NF-κB-α (IκBα), and thereby triggers IκBα degradation. Collectively, these events provoke the nuclear translocation of p65 and p50 subunits of NF-κB, which promotes gene transcription. However, by deubiquitinating TRAF6, deubiquitinases such as the ubiquitin-specific peptidase 20 (USP20) dampen NF-κB activation and subsequent NF-κB-mediated transcription of target genes (11).

Vascular smooth muscle cell (SMC)-specific expression of dominant-negative-USP20 (USP20-DN) increases NF-κB activation, vascular inflammation, and atherosclerotic lesions in *Ldlr*−/− mice (12). Phosphorylation of USP20 on Ser334 regulates the substrate-specific deubiquitinase activity of USP20, in vitro and in vivo (13). In gene-edited mice expressing a USP20 mutant that cannot be phosphorylated on Ser334 (*Usp20*-S334A), arterial injury provokes less SMC NF-κB activation, VCAM-1 expression, and proliferation than it does in WT mice (13). Whether USP20-S334A attenuated neointimal hyperplasia through activity in just SMCs or in SMCs and ECs, too, remains obscure.

The process of angiogenesis, or new blood vessel formation, requires the EC growth factor VEGF-A (VEGF) (14) and its receptor VEGFR2 (15) among other signaling mechanisms. Downstream of VEGFR2 activation in ECs (16) NF-κB activation promotes angiogenesis through mechanisms that include upregulation of EC VEGF (17,18), EC VEGFR2 (1,19), and EC PDGF-BB (1). In addition, EC NF-κB activation promotes EC expression of pro-inflammatory chemokines (20–22) and cell adhesion molecules (16) that attract and retain monocyte/macrophages, which by secreting VEGF also promote angiogenesis (1). Because of the key roles of NF-κB activity in angiogenesis, and because USP20 activity can attenuate NF-κB signaling, we tested the hypothesis that USP20 regulates EC inflammation and angiogenesis.

## Results

### USP20 limits NF-κB activation in ECs

USP20 suppresses cytokine-induced activation of NF-κB in SMCs (11,12). To discern whether USP20 performs a similar role in ECs, we compared aortic primary ECs from *Usp20*^-/-^ and congenic WT mice (Figure 1). As judged by the phosphorylation of NF-κB subunit p65 on Ser536, NF-κB activation in response to IL-1β was substantially greater in ECs from *Usp20*^-/-^ mice than from WT littermate control mice (Figure 1B-C). Thus, USP20 appears to regulate NF-κB activity in ECs, just as it does in SMCs (11–13).

**Figure 1.**
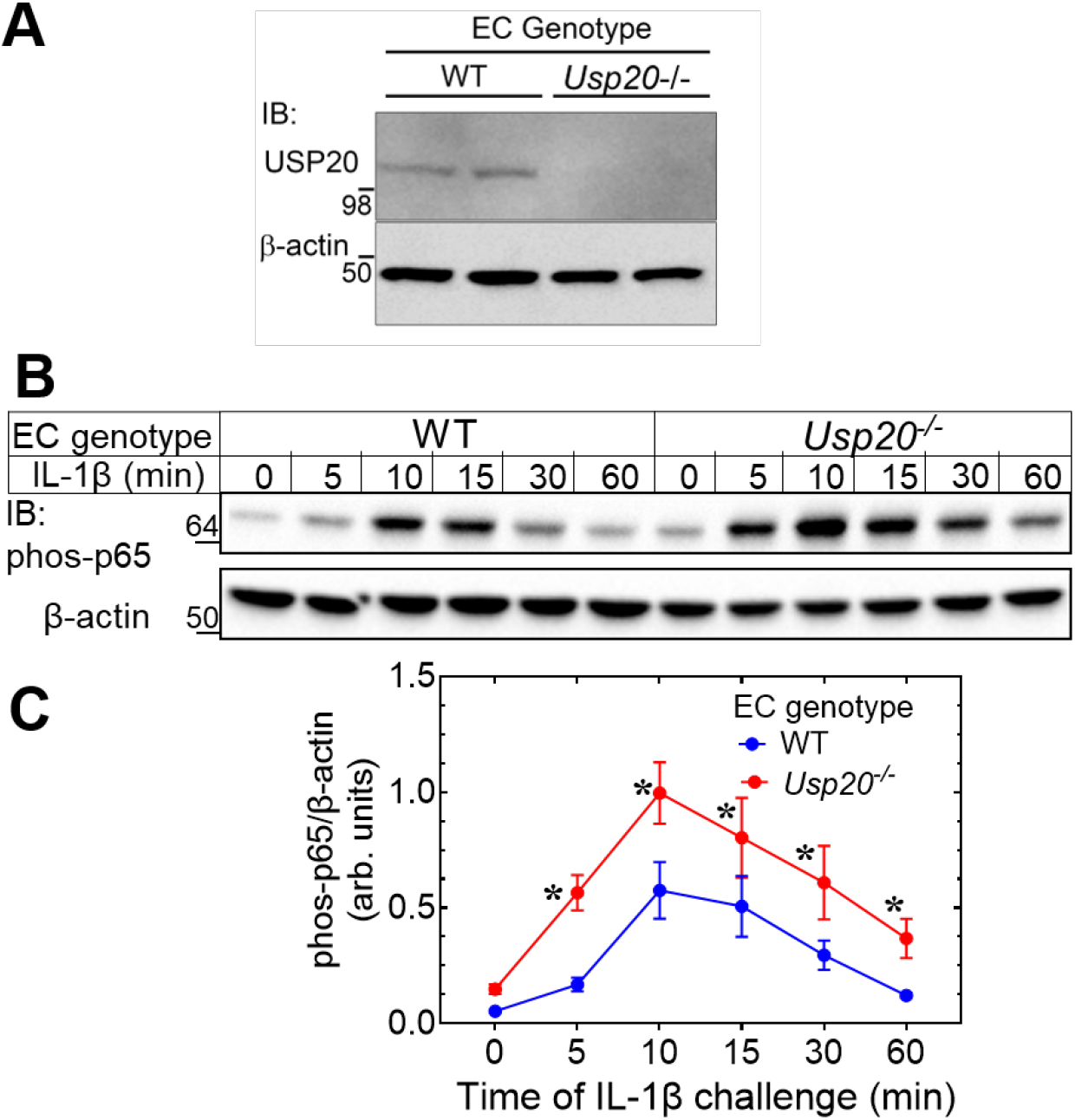
USP20 activity constrains NF-κB activation in primary ECs. (A) Protein extracts of primary aortic ECs isolated from WT and littermate *Usp20*^-/-^ mice were serially immunoblotted to detect USP20 and β-actin. (B) Monolayers of ECs (WT and *Usp20*^-/-^) were serum starved for 4 hours and stimulated with murine IL-1β (10 ng/ml) for the indicated times, and then solubilized. Protein extracts were immunoblotted (IB) serially for the NF-κB p65 subunit phosphorylated on Ser536 (“phos-p65”), and then β-actin. (C) Band densities for phos-p65 were normalized to cognate β-actin band densities within each experiment, and the resulting ratios were plotted as “arbitrary units” (means ± SD) for 3 experiments performed with three independently isolated lines of WT and *Usp20*^-/-^ ECs. Compared with WT: \**p* < 0.01 (2-way ANOVA followed by Holm-Šídák’s multiple comparisons tests).

### USP20 Serine334 phosphorylation increases NF-κB signaling in ECs

The binding of USP20 with its substrate promotes deubiquitinase activity, and this substrate association is regulated by the phosphorylation status of Ser334 or Ser333 in murine and human USP20, respectively (12,13,23,24). Our previous study demonstrated that non-phosphorylated USP20 (USP20-S334A) has increased binding and deubiquitinase activity for TRAF6, and attenuates NF-κB activation more than WT USP20 in SMCs (13). To determine whether USP20 phosphorylation on Ser334 regulates USP20 activity in ECs challenged with inflammatory cytokines, we employed mouse coronary ECs (MCECs) transduced with USP20 constructs in which Ser334 was mutated to mimic (USP20-S334D) or to prevent (USP20-S334A) phosphorylation. IL-1β-induced activation of NF-κB was augmented in MCECs transduced with USP20-S334D, as compared with MCECs transduced with GFP (Figure 2A-B). Indeed, this effect of USP20-S334D was comparable to that of DN-USP20 (Figure 2A-B), consistent with the idea that phosphorylation of USP20 on Ser334 abolished USP20-mediated deubiquitination of NF-κB pathway activators (i.e., TRAF6 (13)). In contrast, IL-1β-induced NF-κB activation was substantially attenuated in MCECs transduced with USP20-S334A or USP20-WT (Figure 2A-B). In MCECs that were not transduced with adenovirus, IL-1β-induced NF-κB activation was similar to that obtained in MCECs transduced with GFP (Figure 2A-B). Thus, in this overexpression system, USP20 activity in ECs negatively regulates IL-1β-induced NF-κB signaling. Furthermore, phosphorylation of USP20 on Ser334 appears to promote NF-κB signaling in ECs in a manner equivalent to the inhibition of USP20 activity.

**Figure 2.**
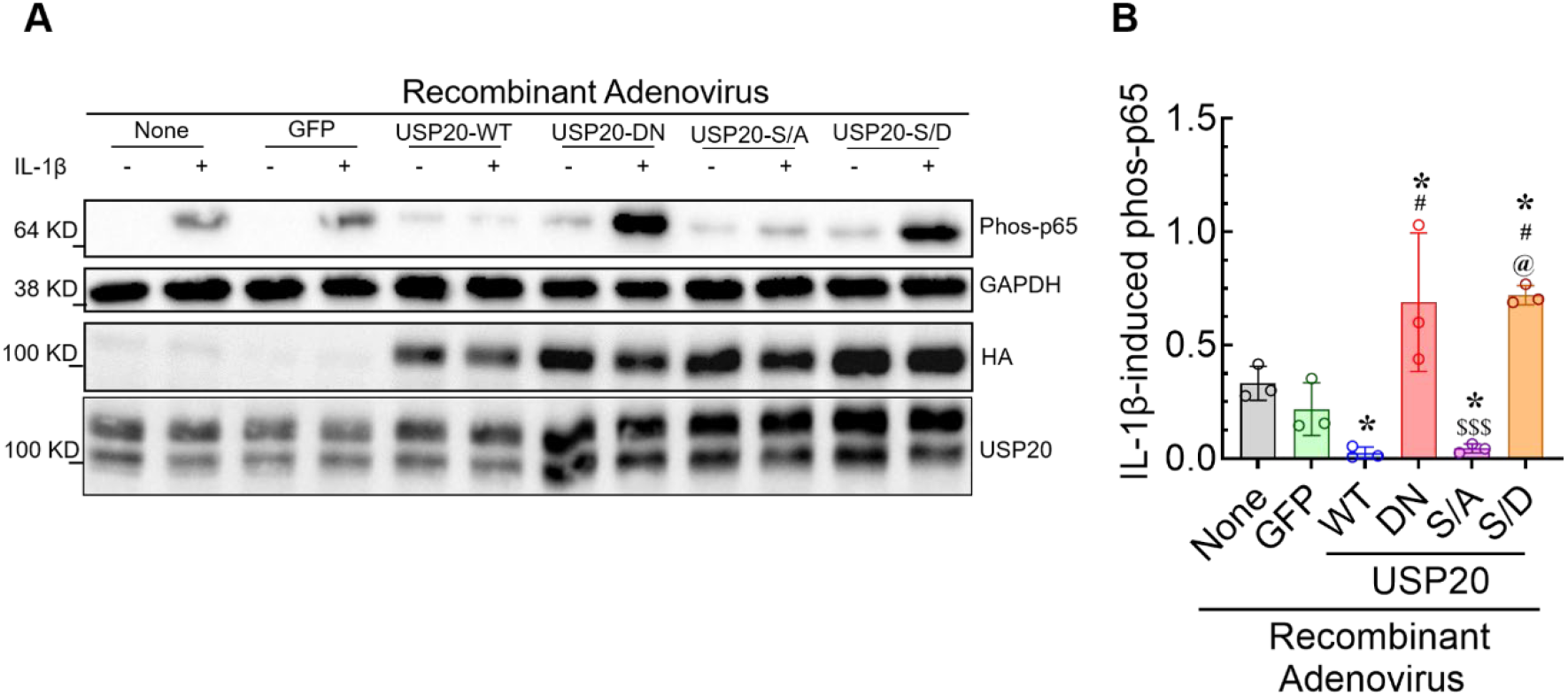
Phosphorylation of USP20 on Ser334 augments NF-κB activation in ECs. (A) MCECs were transduced with no virus (None; control), or with recombinant adenoviruses encoding GFP or HA-tagged isoforms of USP20-WT, USP20-DN, USP20-S334A (S/A), or USP20-S334D (S/D). Cells were then stimulated with (+) or without (−) murine IL-1β (10 ng/mL) for 15 min at 37 °C and lysed. Protein extracts were immunoblotted sequentially for phosphorylated NF-κB p65(Ser536) (“phos-p65”), GAPDH, HA, and USP20. HA immunoblotting confirms expression of recombinant USP20 constructs, while the USP20 blot shows total endogenous and recombinant USP20 protein levels. (B) Densitometric analysis of phospho-p65 band intensities normalized to the corresponding GAPDH band intensities. Data are presented as the mean ± SD from three independent experiments. *, *p* < 0.05 versus control (None); **#,** *p* < 0.05 versus USP20-WT; **$$$,** *p* < 0.001 versus USP20-DN; **@,** *p* < 0.05 versus USP20-S334A (1-way ANOVA followed by Holm–Šídák multiple comparisons test).

### Migration and angiogenic sprouting of ECs are promoted by NF-κB activity

To evaluate the importance of USP20 in regulating NF-κB-dependent angiogenesis, we first examined EC migration, a fundamental process in angiogenesis (25). In an EC monolayer scratch wound-healing assay, migration of MCECs in growth medium was reduced by ∼40% when the NF-κB-activating kinase IKKβ was inhibited selectively with TPCA-1 (Figure 3A-B). In the context of this experiment, it is notable that with 10% fetal bovine serum, the growth medium contains VEGF at ∼35 pM (26), a concentration higher than the K_D_ of the VEGFR2 (∼10 pM (27)). Furthermore, PDGF in fetal bovine serum also activates NF-κB in ECs—either through the PDGF receptors themselves (28) or through the VEGFR2, to which PDGF-BB binds with relatively high affinity (KD ∼40 nM (27)). To test capillary-like tube formation, we used an EC spheroid angiogenesis (sprouting) assay (29) driven by VEGF at approximately 750 pM (VEGFR2-saturating (27)). MCEC sprouting was reduced by ∼70% in TPCA-treated MCECs, as compared with vehicle-treated MCECs (Figure 3C-D). Together, these results strongly suggest that NF-κB activation is required for EC migration and VEGF-driven spheroid outgrowth/angiogenesis in these model systems.

**Figure 3.**
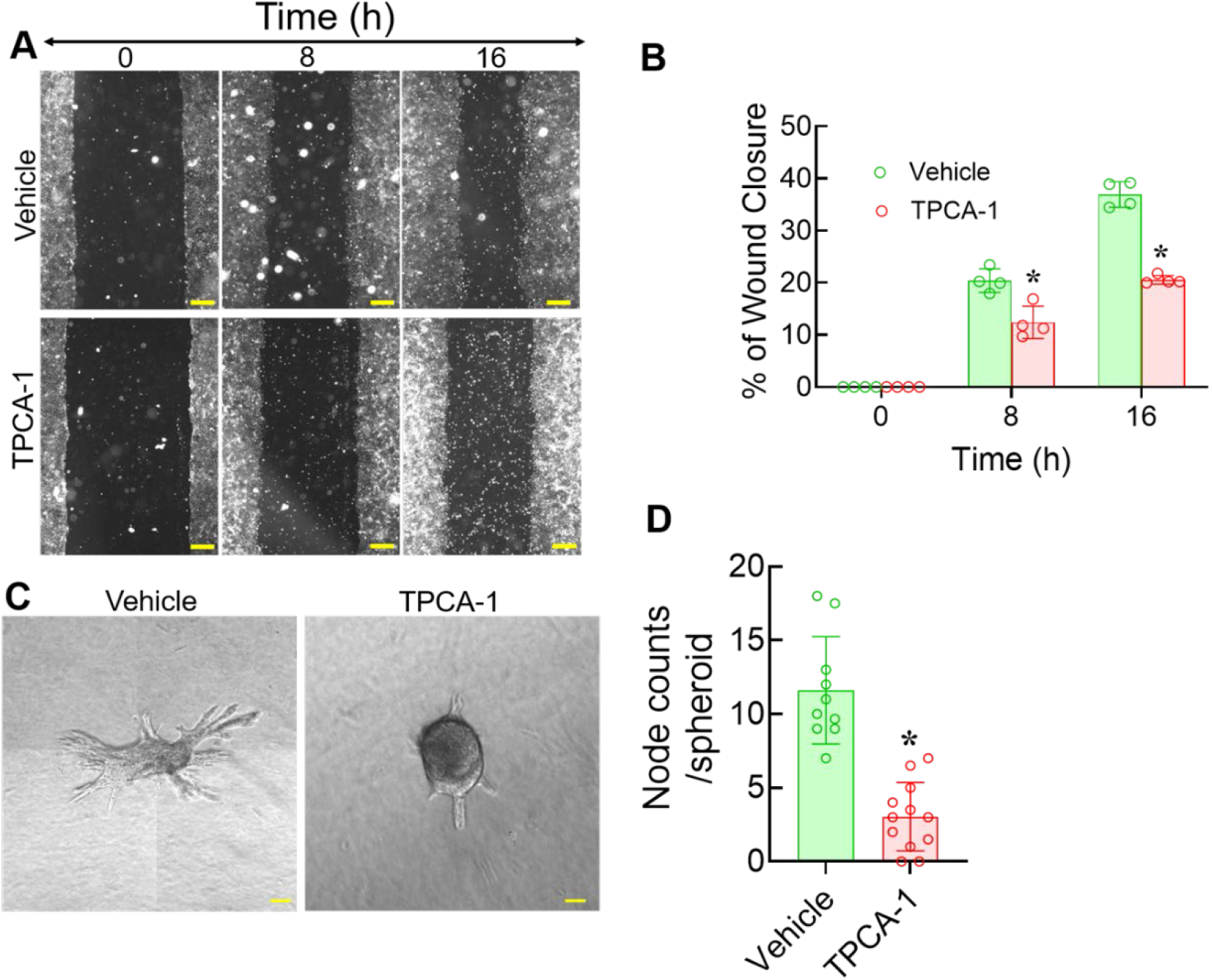
Migration and sprouting of MCECs are blocked by TPCA-1. (A) Confluent monolayers of MCECs were subjected to scratch-wound assays in the absence (vehicle) or presence of the IKKβ inhibitor TPCA (10 μM), and photographed with a phase-contrast microscope at the indicated times after the scratch wound to the monolayer. Scale bar = 200 µm. (B) Wound closure was quantified as described in the methods from 4 independent experiments and at least 6 images for each condition and the average wound area ± SD is summarized in the graph. *, *p* < 0.01 vs vehicle as indicated, two-way ANOVA followed by Holm-Šídák’s multiple comparison tests. (C) MCEC spheroids were embedded in type I collagen gel and incubated with 30 ng/mL VEGF in the presence of vehicle or 10 µM TPCA-1 for 48 hr (37 °C). Representative photomicrographs are shown. Scale bars = 20 µm. (D) The number of EC sprout branch points (nodes) were counted by observers blind to specimen identity, and plotted (with means ± SD) for 2-3 independent spheroids of each treatment group. *, *p<*0.01 vs vehicle control (unpaired t-test).

### USP20 activity diminishes EC migration and angiogenic sprouting

Because USP20 activity in ECs diminishes NF-κB activation (Figs 1, 2), and because NF-κB activation appears necessary for EC migration and angiogenic sprouting (Fig 3), we tested whether USP20 could inhibit EC migration and angiogenic sprouting. To that end, we performed the EC monolayer scratch-wound healing assay using MCECs transduced with recombinant adenoviruses encoding GFP, USP20-WT, USP20-DN, USP20-S334A, or USP20-S334D. Compared with control MCECs transduced with GFP, MCECs transduced with either USP20-DN or USP20-S334D demonstrated enhanced migration, whereas MCECs transduced with either USP20-WT or USP20-S334A demonstrated diminished migration (Figure 4A-B). The expression of different USP20 variants was equivalent by immunoblotting (Figure 2, HA immunoblot). The transduction efficiency of all the recombinant adenoviruses was comparable, with the exception of USP20-WT, for which the prevalence of bright-staining MCECs was <50% by immunostaining and confocal microscopy (Figure 4C). Thus, in ECs USP20 activity inhibits both NF-κB activation (Figures 1-2) and EC migration.

**Figure 4.**
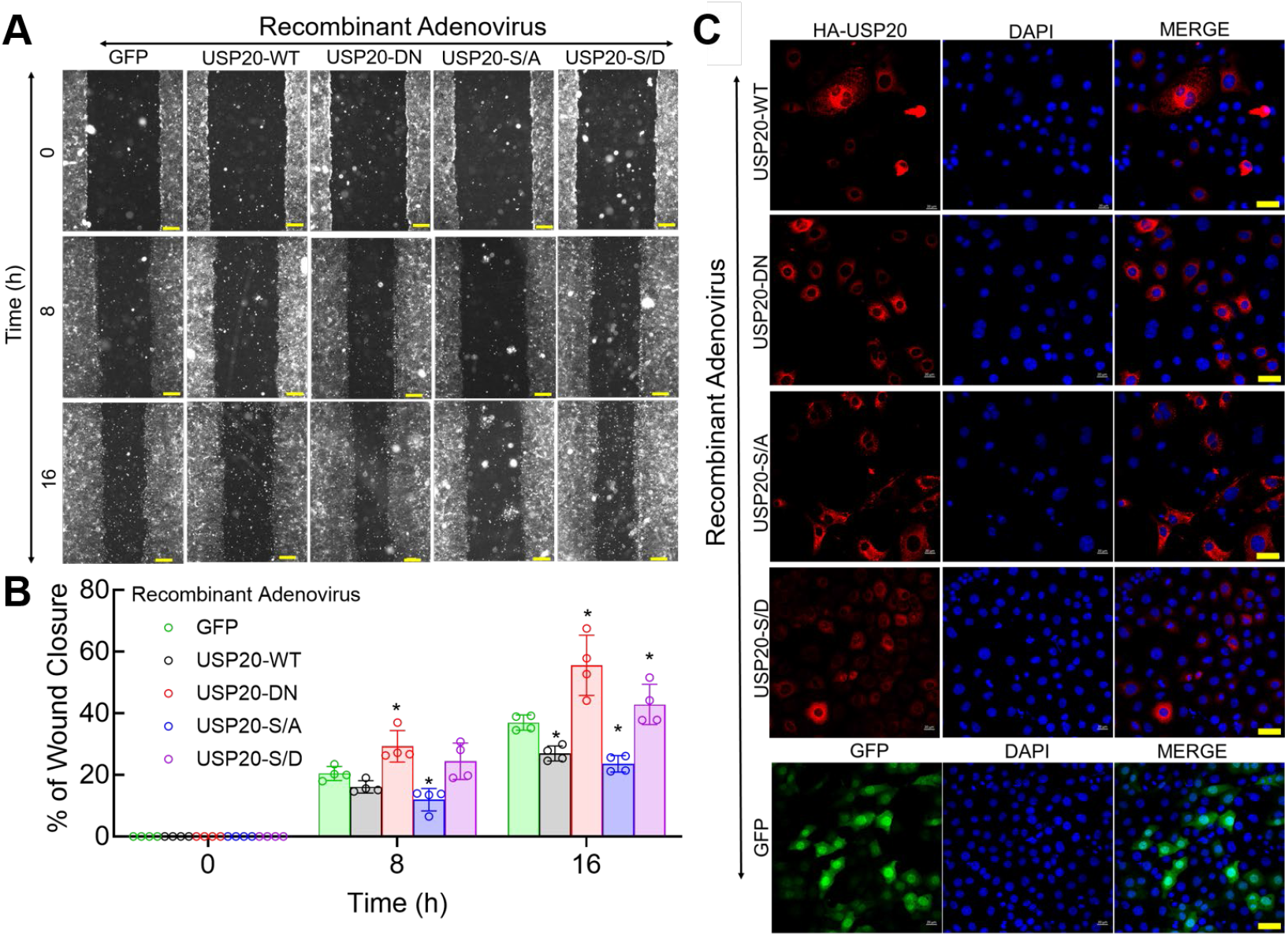
USP20 activity constrains EC migration. (A) MCECs were transduced with recombinant adenoviruses encoding GFP (control) or the N-terminal HA-tagged USP20 isoforms used in Fig 2. Forty-eight hours after transduction, confluent monolayers were wounded and analyzed for migration as in Fig 3. Scale bar = 200 μm. (B) EC migration was quantified as in Fig 3, and presented as mean ± SD from four independent experiments. Compared with GFP: *, *p* < 0.05 (2-way ANOVA followed by Holm–Šídák’s multiple comparisons test). (C) MCECs were transduced with recombinant adenoviruses expressing GFP or the indicated HA-tagged USP20 constructs. HA-tagged USP20 was detected by anti-HA immunostaining (red); GFP detection is shown in green, and nuclei were counterstained with DAPI (blue). Scale bar = 20 μm.

To test the role of USP20 and its phosphorylation status in EC sprouting, we performed the spheroid-based sprouting assay (Figure 5). Concordant with our MCEC migration data (Figure 4), transduction of active and inactive USP20 constructs produced reciprocal results. Whereas USP20-WT and USP20-S334A overexpression reduced the MCEC spheroid sprouting observed in control spheroids, USP20-DN and USP20-S334D overexpression augmented MCEC spheroid sprouting (Figure 5). Thus, USP20 phosphorylation status and catalytic activity appear to regulate EC sprouting in a manner that parallels the modulation of NF-κB activity by these USP20 species.

**Figure 5.**
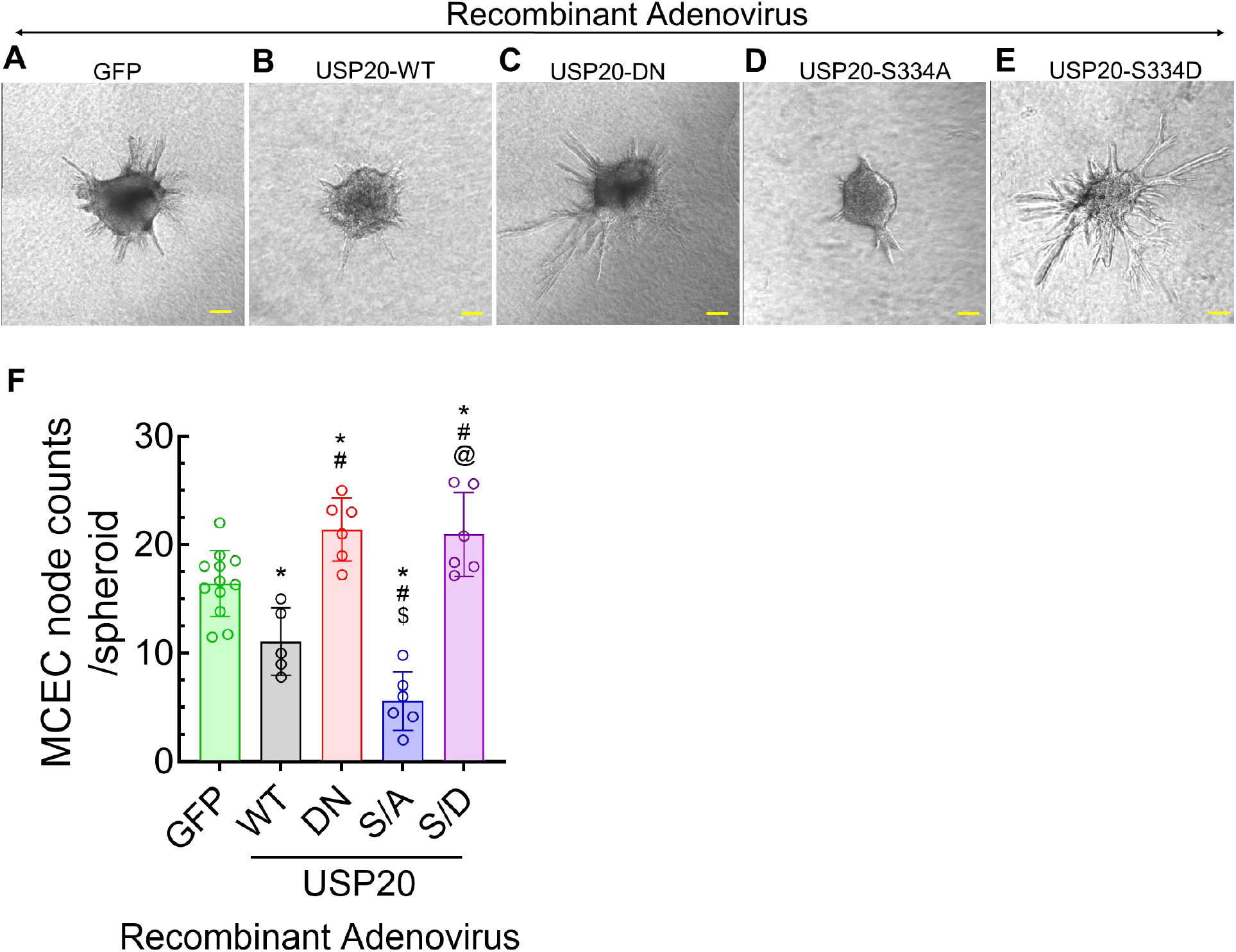
USP20 activity diminishes angiogenic sprouting of EC spheroids. (A-E) MCECs were transduced with recombinant adenoviruses encoding GFP or the indicated USP20 isoform as in Figs 2 and 4. Twenty-four hours after transduction, approximately 400 cells were used to generate spheroids by the hanging-drop method. Spheroids were embedded in a collagen type I matrix and stimulated with VEGF (30 ng/mL) for 48 hours to induce angiogenic sprouting. Representative phase-contrast micrographs from one of five independent experiments are shown. Scale bars = 20 µm. (F) Angiogenic sprouting was quantified by determining the number of sprout branch points (nodes) per spheroid using ImageJ. Data are presented as means ± SD from 5–12 wells, with each well containing 1–5 spheroids. Compared with GFP (*), USP20-WT (#), USP20-DN ($), or USP20 S334A (@): *p* < 0.05 (one-way ANOVA and Holm–Šídák’s multiple comparisons test).

To discern whether our findings with USP20 overexpression in vitro also obtained with physiologically expressed USP20, we used VEGF to stimulate EC sprouting from aortic rings (30–32) excised from *Usp20*^-/-^ and WT mice. EC sprouting was significantly increased from aortic rings of *Usp20*^-/-^ mice, as compared with aortic rings of WT mice (Figure 6A-C, S3A-C). Thus, physiologically expressed USP20 activity appears to constrain angiogenesis assessed as aortic sprouting.

**Figure 6.**
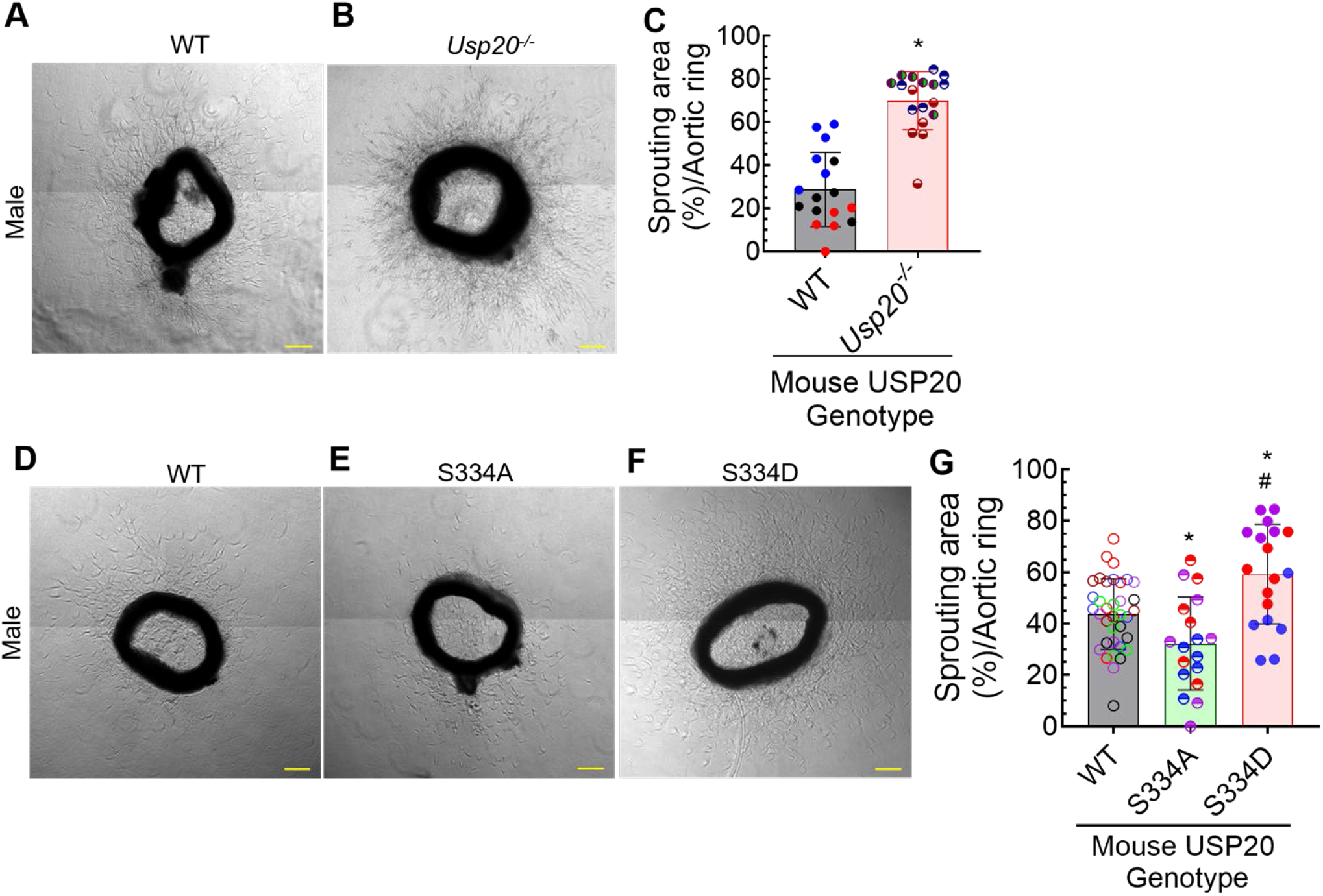
USP20 regulates angiogenesis in a phosphorylation-dependent manner. (A–B) Thoracic and abdominal aortic sections isolated from 3-month-old male WT and *Usp20*^-/-^ mice were cut into approximately 1-mm rings, embedded in collagen type I gel, and cultured in endothelial growth medium supplemented with VEGF (30 ng/mL) to stimulate microvessel sprouting. Representative phase-contrast images acquired after 5 d of culture are shown. Scale bar = 200 μm. (C) Angiogenic sprouting was quantified by measuring the total sprouting area surrounding each aortic ring using ImageJ. Data are presented as mean ± SD from three mice per genotype, with multiple aortic rings analyzed from each mouse. Compared with WT: *, *p* < 0.01 (t-test). (D–F) Aortic ring microvessel sprouting was performed as in panel A, but with aortic rings from 3-month-old male WT, *Usp20*-S334A (phospho-resistant), and *Usp20*-S334D (phospho-mimetic) mice. Representative phase-contrast images obtained after 5 days of VEGF stimulation are shown. Scale bar = 200 μm. (G) Sprouting area was quantified as in panel C. Data are presented as mean ± SD from three mice per genotype, with multiple aortic rings analyzed from each mouse. Compared with WT: *, *p* < 0.05; compared with USP20-S334A: #, *p* < 0.01 (1-way ANOVA followed by Holm–Šídák’s multiple comparisons test). Each mouse represents 6 dots of the same color in each bar; each dot represents the average sprouting area of 5-7 aortic rings.

To test further whether angiogenesis depends upon USP20 phosphorylation, we employed aortic rings from congenic, gene-edited mice expressing physiologic levels of mutated USP20: *Usp20*-S334A or *Usp20*-S334D. As we have reported previously (13) and shown in Figures 2-4, phosphorylation of USP20 on Ser334—mimicked by the S334D mutation—diminishes USP20 activity on TRAF6, as well as on other substrates (23). In accord with these data, aortic ring EC sprouting was significantly greater from *Usp20*-S334D than from WT aortas (Figure 6 D-G, Figure S3 D-G). In contrast, the aortic ring EC sprouting was significantly less from *Usp20*-S334A than from WT aortas (Figure 6 D-G, Figure S3, D-G).

### NF-κB inhibitor diminishes sprouting of aortic rings prepared from WT and Usp20⁻/⁻ mice

EC NF-κB signaling (Figure 1) and aortic EC sprouting (Figure 6 A-C) are both increased by USP20 deficiency. To determine whether augmented EC sprouting from *Usp20*^-/-^ aortic rings involves augmented NF-κB activity, we employed aortic rings of WT and *Usp20*^-/-^ mice to perform the aortic ring EC sprouting assay in the presence of the IKKβ inhibitor TPCA-1. As shown in Figure 7, inhibition of NF-κB activity with TPCA-1 decreased aortic ring sprouting in WT and *Usp20*^-/-^ mice by 44-81% compared with their respective controls (Figure 7A-D)—in aortic rings taken from both male (Figure 7A-B) and female (Figure 7C-D) mice. These results suggest that EC sprouting from aortic rings of both WT and *Usp20*^-/-^ mice requires NF-κB activation.

**Figure 7.**
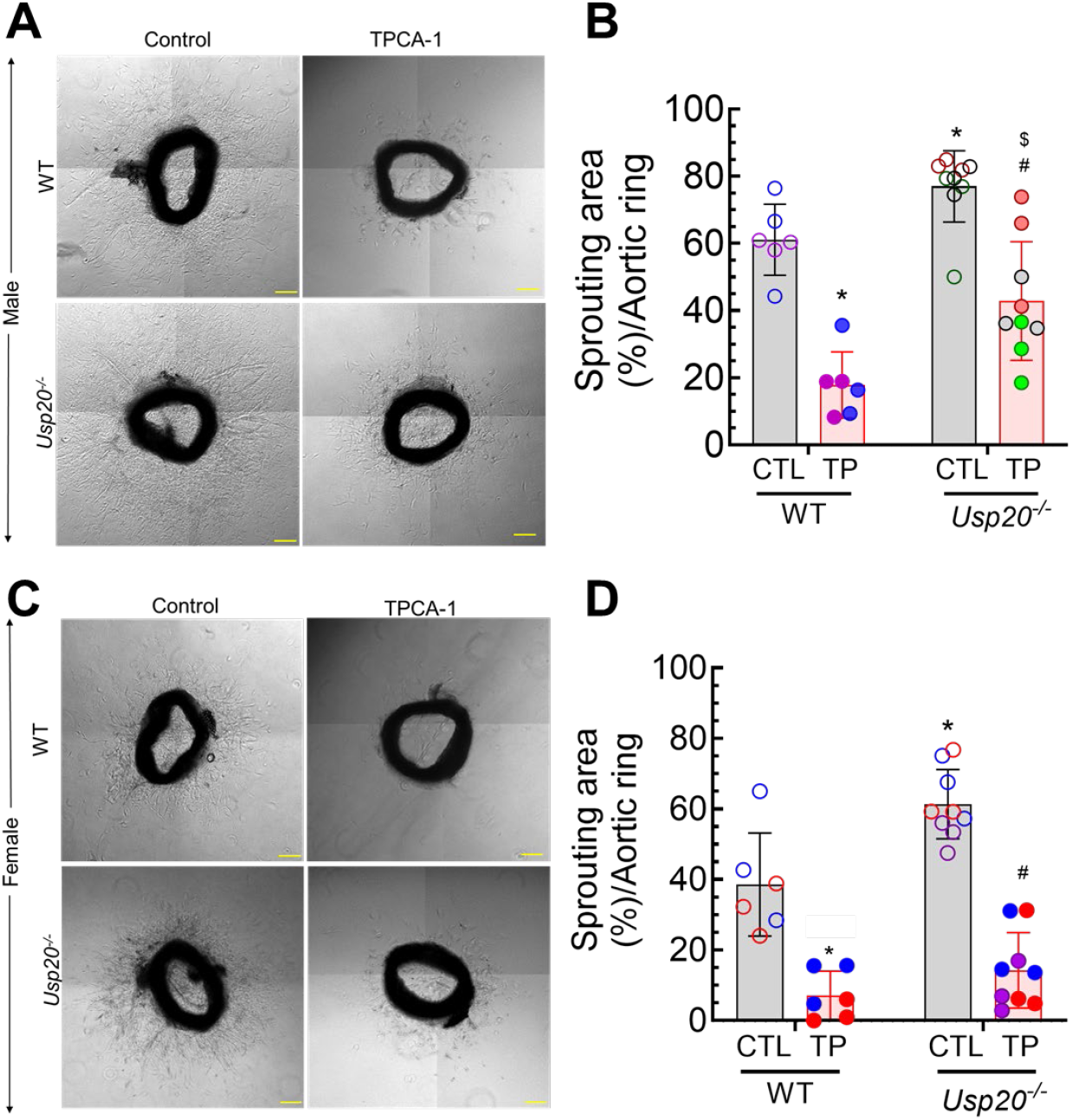
NF-κB inhibition decreases angiogenesis. Aortic ring EC sprouting assays were performed with aortic rings from male WT and *Usp20*^-/-^ mice as in Fig 6A-C, except that rings were treated with vehicle lacking (control, CTL) or containing the IKKβ inhibitor TPCA-1 (10μM) for 5 days. (A) Representative phase-contrast images. Scale bars = 200 μm. (B) Quantification of sprouting area. Data are mean ± SD. Aortic rings were obtained from 2 WT and 3 *Usp20*^-/-^ mice. *, *p* < 0.01 vs WT control; #, *p* < 0.01 vs *Usp20*^-/-^ control; $, *p* < 0.01 vs WT + TPCA-1, as indicated (two-way ANOVA with Holm-Šídák post hoc test). (C) Representative images of VEGF-stimulated aortic ring sprouting from 3-month-old female WT and *Usp20*^-/-^ mice cultured under the same conditions as in (A). Scale bar = 200 μm. (D) Quantification of sprouting area. Data are mean ± SD. Aortic rings were obtained from 2 WT and 3 *Usp20*^-/-^ mice. *, *p* < 0.01 vs WT control; #, *p* < 0.01 vs *Usp20*^-/-^ control, as indicated (two-way ANOVA with Holm-Šídák post hoc test). In panels B and D, each mouse is represented by 3 dots of the same color in each bar, with each dot representing the average sprouting area of 5–7 aortic rings. Abbreviations: WT, wild type; TPCA-1 (TP), IKKβ/NF-κB inhibitor.

### Identification of angiogenic factors modulated by USP20 activity

To identify angiogenic factors whose expression may be regulated by USP20, we solubilized EC sprouts from WT and *Usp20*⁻/⁻ aortic rings to extract EC proteins, and assayed these extracts on a Proteome Profiler Mouse Angiogenesis Array (R&D Systems). Among the proteins upregulated in *Usp20*⁻/⁻ sprouts relative to WT (Figure 8A-C), several are established transcriptional targets of NF-κB: MMP-3 (33,34), PAI-1 (35), VEGF (36,37), and TIMP-1 (38). Additional proteins identified on the array — including CXCL12 and IGFBP-9 — are shown in Figure 8B-C, but their regulation by NF-κB is less well established. Because USP20 suppresses NF-κB activation in ECs (Figure 1), the increased expression of these NF-κB-dependent pro-angiogenic factors in *Usp20*⁻/⁻ sprouts directly supports the conclusion that USP20 constrains EC angiogenic signaling by limiting NF-κB activation. Among the upregulated proteins, MMP-3 was selected for further investigation because it exhibited one of the largest increases in expression and is a known NF-κB-regulated gene (33,34).

**Figure 8.**
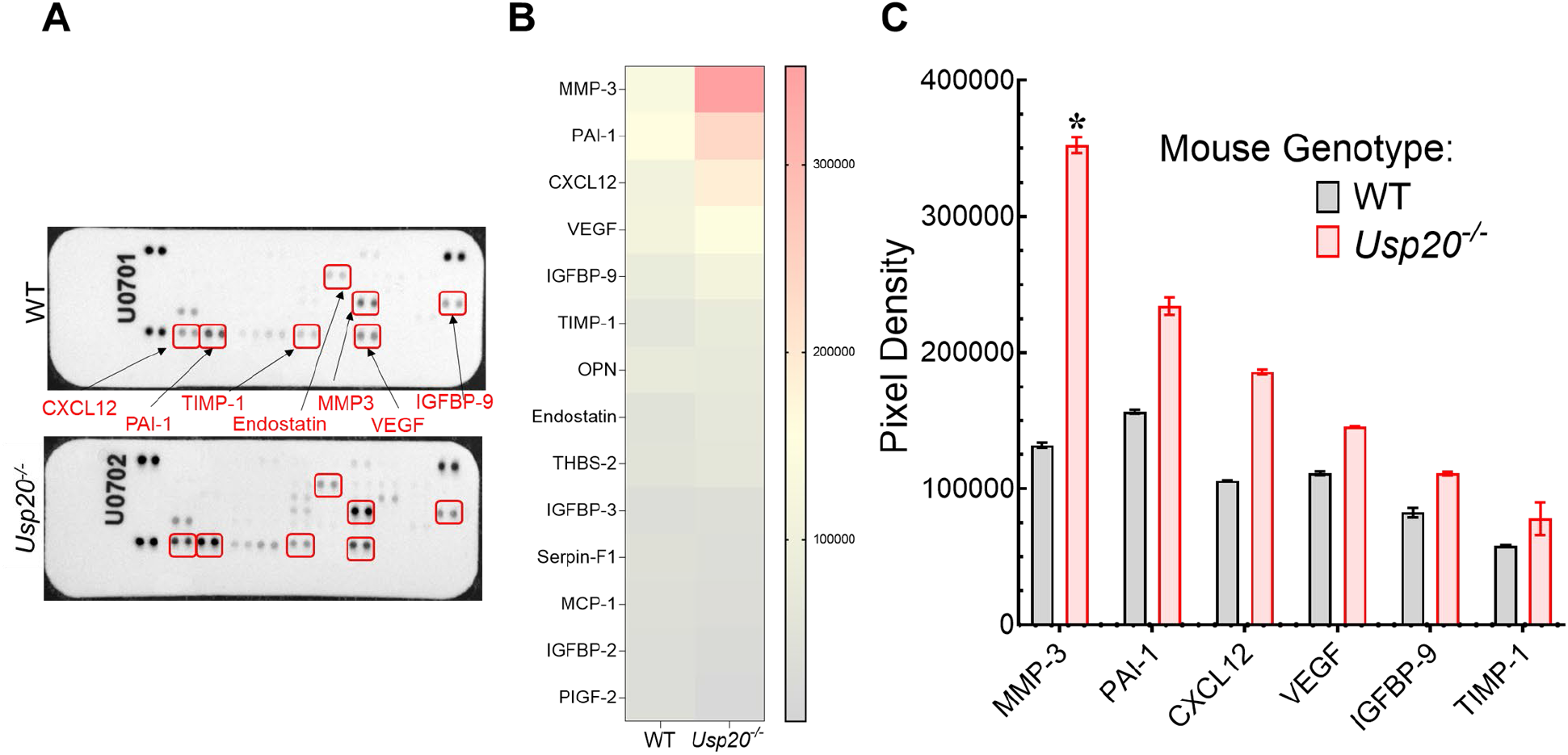
Proteomic profiling identifies angiogenic proteins upregulated in *Usp20*−/− aortic sprouts. (A) Thoracic and abdominal aortic rings from 3-month-old WT and *Usp20*^-/-^ mice were cultured in collagen type I gels in the presence of VEGF (30 ng/mL) for 5 days to induce angiogenic sprouting. Sprouting tissues were harvested, lysed, and equal amounts of total protein were analyzed using a mouse angiogenesis antibody array containing 53 angiogenesis-related proteins. Representative array membranes are shown, with selected differentially expressed proteins indicated. (B) Densitometric analysis of angiogenesis-related proteins following normalization to internal positive-control spots on each membrane. The heatmap displays the 15 proteins showing the largest differences in expression between *Usp20*^-/-^ and WT aortic sprouts. (C) Quantification of selected proteins exhibiting increased expression in *Usp20*^-/-^ aortic sprouts relative to WT. Data are presented as normalized signal intensity (mean ± SD) from two independent biological samples per genotype. Statistical significance was assessed using an unpaired two-tailed Student’s t-test. \**p* < 0.05 versus WT.

### MMP-3 promotes angiogenic sprouting in Usp20⁻/⁻ mice

Since MMP-3 was increased in sprouts from *Usp20*⁻/⁻ aortic rings (Figure 8), and endothelial MMP-3 expression is upregulated by NF-κB (33), we further focused on this protein. To test whether MMP-3 contributes to the augmented angiogenesis observed in *Usp20*⁻/⁻ aortic rings, we assayed VEGF-stimulated aortic rings from *Usp20*⁻/⁻ mice in the presence and absence of MMP-3 inhibitor II (39). MMP-3 inhibitor II significantly reduced angiogenic sprouting from *Usp20*⁻/⁻ aortic rings by ∼47% (Figure 9), demonstrating that MMP-3 contributes functionally to the excess sprouting observed in the absence of USP20. These findings suggest that USP20 restrains angiogenesis in part by attenuating NF-κB-driven MMP-3 expression.

**Figure 9.**
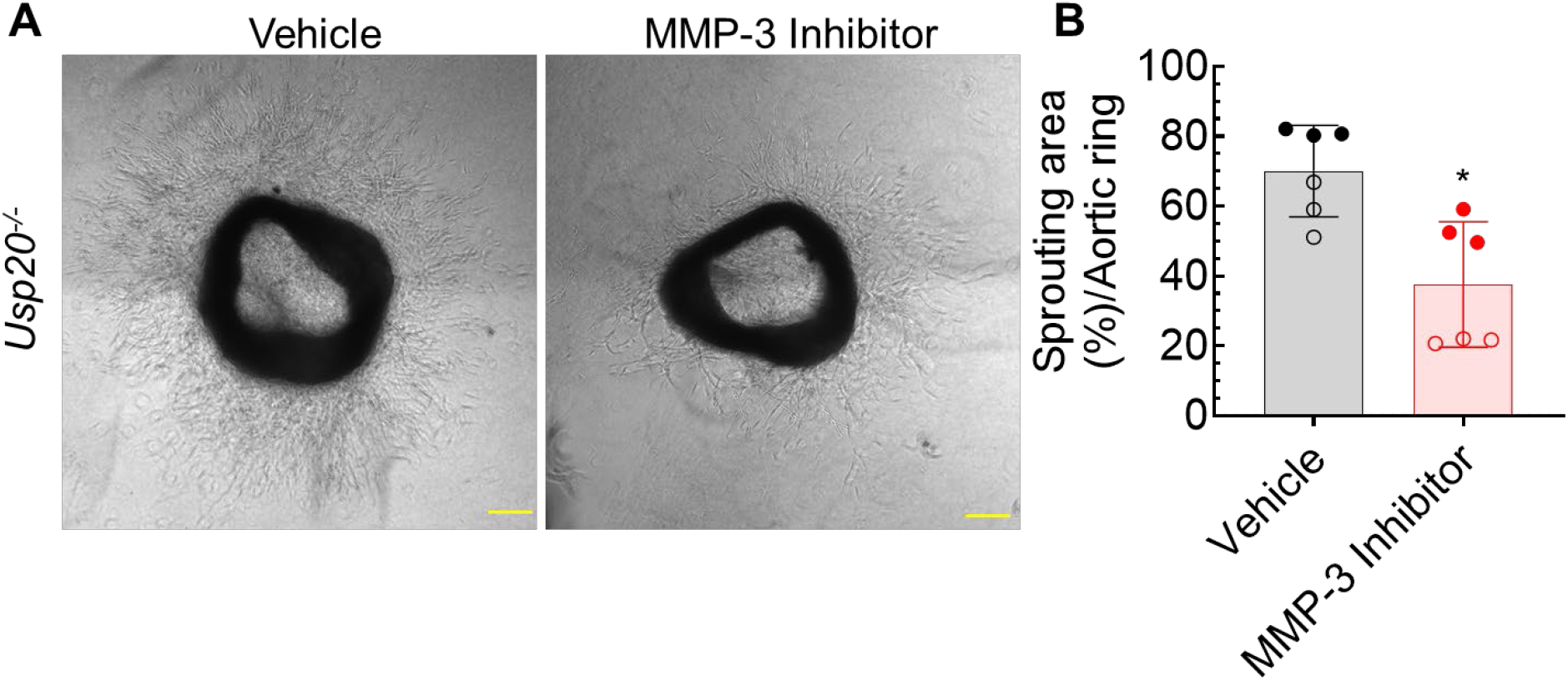
Aortic sprouting in *Usp20^-/-^*samples is attenuated by MMP-3 inhibition. (A) Thoracic and abdominal aortic rings were isolated from 3-month-old male *Usp20*^-/-^ mice as in Fig 6, in the absence (vehicle) or presence of MMP-3 inhibitor II (1 μM). Representative phase-contrast images of microvessel sprouting were acquired after 5 days of culture. Scale bar = 200 μm. (B) Angiogenic sprouting was quantified by measuring the total sprouting area surrounding each aortic ring using ImageJ. Data are presented as mean ± SD. Aortic rings were obtained from 2 *Usp20*^-/-^ male mice. Each mouse is represented by six symbols (three vehicle-treated and three MMP-3 inhibitor II-treated), with each symbol representing the average sprouting area of seven aortic rings. Compared with vehicle-treated: *, *p* < 0.05 (t-test).

## Discussion

Our findings demonstrate a novel role for USP20 in constraining EC inflammatory signaling, sprouting, and angiogenic growth. USP20 diminishes ubiquitination-dependent NF-κB activation elicited by IL-1β and, in the context of VEGF-A-stimulated sprouting, appears to attenuate NF-κB-dependent angiogenic signaling in a manner that is regulated by phosphorylation of USP20 on Ser334. By inhibiting NF-κB signaling, USP20 activity diminishes the expression of MMP-3 and other pro-angiogenic factors. USP20’s anti-inflammatory activity inversely correlates with Ser334 phosphorylation, which increases in both SMCs and ECs with atherogenesis (13), suggesting that inhibiting USP20 Ser334 phosphorylation could be a therapeutic target in atherosclerosis (40) and other NF-κB-driven diseases (5,41–43).

Our prior work established that USP20 deubiquitinates TRAF6 (11,12) and RIPK1 (12) in SMCs, attenuating SMC NF-κB signaling and atherosclerosis (11–13). The current study extends this paradigm to ECs: USP20 suppresses IL-1β-induced NF-κB activation in primary ECs and MCECs (Figures 1–2), and USP20 deficiency or S334D expression augments VEGF-A-driven aortic ring sprouting (Figure 6). *Usp20*-S334A mice also developed ∼50% less neointimal hyperplasia than WT mice after carotid endothelial denudation (13); the present finding that *Usp20*-S334A suppresses aortic ring angiogenesis extends this protection to the endothelial compartment, reinforcing Ser334 phosphorylation as a therapeutic target.

Phosphorylation of USP20 Ser334 by IRAK1 diminishes the association of USP20 with its substrates, reducing its deubiquitinase activity toward TRAF6 without altering intrinsic catalytic activity per se (13,23). This substrate-disengagement mechanism explains a key observation in our data: overexpressed USP20-S334D behaves like the dominant-negative USP20 construct (USP20-DN), which lacks catalytic activity, because both fail to engage and deubiquitinate TRAF6 and thence activate NF-κB (Figure 2). When expressed at physiological levels in the *Usp20*-S334D CRISPR mouse, perhaps a retention of partial substrate engagement occurs— consistent with the intermediate aortic ring sprouting phenotype we observed: greater than WT but less than *Usp20*^-/-^ (Figure 6 D–G). Conversely, the phospho-resistant *Usp20*-S334A mutant maintains constitutive substrate engagement, explaining its attenuated NF-κB signaling and angiogenic sprouting. Because USP20 Ser334 phosphorylation is elevated in SMCs from human atherosclerotic segments, (13) a parallel elevation in ECs would constitute a pathological amplification loop: IRAK1 phosphorylates USP20-Ser334 → TRAF6 accumulates K63-ubiquitin → NF-κB drives cytokine production and angiogenic gene expression. Strategies that prevent this phosphorylation could thus simultaneously constrain EC neovascularization and SMC-driven inflammation — a dual benefit relevant to atherosclerosis and restenosis.

In our aortic ring sprouting assay, VEGF-A serves as the primary angiogenic stimulus, yet USP20 activity constrains this sprouting response. Our data do not establish the precise upstream mechanism by which TRAF6-dependent NF-κB signaling is engaged during angiogenic sprouting. However, multiple inflammatory and angiogenic pathways converge on NF-κB signaling in endothelial cells (1,2,5), and inhibition of NF-κB with TPCA-1 markedly suppressed sprouting in both spheroid and aortic ring assays (Figures 3 and 7). Consistent with our previous studies demonstrating that USP20 deubiquitinates TRAF6 and attenuates NF-κB signaling (11–13), the present findings support a model in which USP20 constrains a shared ubiquitin-dependent NF-κB signaling node required for endothelial migration and angiogenic sprouting. Augmented sprouting when USP20 activity is impaired (*Usp20*−/−, or *Usp20*-S334D; Figures 5 and 6) and reduced sprouting when USP20 activity is enhanced (*Usp20*-S334A; Figures 5 and 6) further support the conclusion that USP20 negatively regulates endothelial angiogenic responses through suppression of NF-κB signaling. Future studies will be required to identify the upstream inflammatory or angiogenic signals that engage TRAF6-dependent NF-κB signaling during endothelial sprouting.

Our three experimental systems employ distinct proximal stimuli — IL-1β (NF-κB western blots), VEGF-A (aortic ring and spheroid assays), and FBS (scratch-wound assay) — yet TPCA-1 suppressed NF-κB-dependent responses in all three (Figures 3 and 7), suggesting a shared IKK→NF-κB node. IL-1β activates this node via IL-1R →IRAK1→TRAF6→IKK (44,45). FBS contains VEGF-A (14–21 ng/mL) (26) and IL-1 family cytokines that independently stimulate VEGF synthesis via VEGFR-2, (46,47) and proteomes of IL-1β- and VEGF-A-stimulated HUVECs show substantial overlap in inflammatory and angiogenic proteins (48). Together, these observations suggest that, regardless of the proximal trigger, a shared ubiquitin-dependent NF-κB node is engaged — and that USP20, by deubiquitinating TRAF6, can attenuate this node whether the initiating signal is inflammatory or angiogenic.

Our proteome profiler analyses identified MMP-3 (49) as a candidate NF-κB-regulated angiogenic factor whose expression is increased in *Usp20*^-/-^ sprouts. Absence of USP20 engenders a marked increase in NF-κB signaling and MMP-3 expression in aortic angiogenesis. Inhibition of MMP-3 with MMP-3 Inhibitor II significantly decreased angiogenic sprouting in *Usp20*⁻/⁻ aortic rings (∼47%; Figure 9). Inflammation-induced upregulation of MMP-3 is mediated via NF-κB activation in vascular SMCs, (50) and NF-κB directly regulates the MMP-3 promoter through binding of its p50 and p65 subunits (49). Our aortic ring assay embeds rings in a collagen type I matrix, yet MMP-3 does not cleave fibrillar collagen I — its substrates include collagens III, IV, V, IX, X, and XI, fibronectin, laminin, and proteoglycans, but not fibrillar collagen I (49). How then does MMP-3 Inhibitor II attenuate sprouting in our collagen I matrix? MMP-3 is well established as an activator of pro-MMP-1 and pro-MMP-9, (49,51) and both MMP-1 and MMP-9 cleave fibrillar collagen I, thereby enabling EC migration through the matrix (49). We therefore propose that elevated MMP-3 (in *Usp20*^-/-^ or *Usp20*-S334D conditions) promotes angiogenesis indirectly: MMP-3 activates pro-MMP-1 and pro-MMP-9 — the former being the principal interstitial collagenase responsible for cleaving fibrillar collagen I — which then degrade the scaffold and enable EC sprouting. MMP-3 Inhibitor II, by blocking this cascade, would therefore impair EC migration regardless of the collagen type present. MMP-3 is a prognostic marker of atherosclerotic plaque vulnerability and ischemic heart disease (49,52); our data now link it functionally to USP20-regulated, NF-κB-driven angiogenesis in the vascular endothelium.

Taken together, our studies reveal that EC USP20 negatively regulates angiogenesis by attenuating ubiquitination-dependent NF-κB signaling and its downstream effector MMP-3. The convergence of IL-1β, TNF, and VEGF-A signaling on a TRAF6-dependent ubiquitin–IKK–NF-κB axis places USP20 at a nodal position capable of modulating EC responses across multiple inflammatory and angiogenic stimuli. Molecules or compounds that activate USP20, or that prevent its Ser334 phosphorylation, therefore represent potentially novel therapeutic candidates to mitigate pathological angiogenesis in atherosclerotic plaque growth and solid tumor malignancies.

## Experimental Procedures

### Materials

Murine IL-1β was purchased from PeproTech. Recombinant mouse VEGF (Cat. No. 493-MV) was from R&D Systems; methylcellulose (Cat. No. M7027) and rat collagen type I (Cat. No. 08-115) were from Millipore Sigma; Opti-MEM (Cat. No. 51985–034) was from Thermo Fisher Scientific; Antibodies used for immunoblotting and immunofluorescence are listed below, along with their sources. All antibodies were diluted at 1:1000 for immunoblotting and 1:100 for immunofluorescence, unless stated otherwise. Rabbit monoclonal anti-phospho-p65(Ser536) (Cat. No. 3033, 1:5000), and rabbit monoclonal anti-GAPDH conjugated to HRP (Cat. No. 3683S, 1:5000) were from Cell Signaling Technologies; mouse monoclonal anti-β-actin (Cat. No. A5441, 1:10,000) was from Millipore Sigma. Mouse monoclonal anti-CD31 (Cat. No. MA5-13188) and Alexa Fluor 594-conjugated wheat germ agglutinin (WGA; Cat. No. W11262) were from Thermo Fisher Scientific; Horseradish peroxidase-conjugated secondary antibodies were from Cell Signaling Technology, Rockland Immunochemicals and Bethyl Laboratories, Inc; they were used at a dilution of 1:5000. Secondary antibodies conjugated to Alexa Fluor 488® and Alexa Fluor 546® were obtained from Invitrogen; they were used at 1:100. MMP-3 inhibitor II (Cat. No. 444225) was from Millipore Sigma.

### Mice

All animal experiments were conducted in accordance to protocols that were approved by the Duke University Institutional Animal Care and Use Committee. From the Sanger Institute, we obtained *Usp20* mice with a genetrap null allele (Usp20tm1a(EUCOMM)). Mating this with Flp-transgenic mice (strain # 009086, Jackson Labs) removed the lacZ and neo cassettes of the “knockout-first” allele, (53) thereby restoring *Usp20* expression and rendering the *Usp20* allele “floxed” (loxP sites flank exons 7 and 8). By breeding *Usp20*^flox/flox^ with CMV-Cre mice, we obtained *Usp20*^-/-^ (USP20-KO) mice, which are born at expected rates and are phenotypically normal under non-stressed conditions. *Usp20*^-/-^ mice were back-crossed into C57BL/6J, and heterozygous *Usp20*^-/+^ were bred to obtain *Usp20*^+/+^ wild type (WT) and *Usp20*^-/-^ littermates that were used in this study (Figure S1). *Usp20*-S334D and *Usp20*-S334A mice were generated by the CRISPR/Cas9-mediated gene editing with the help of Duke Transgenic Core facility (Figure S2).

### Cell lines

Aortic ECs were isolated from the descending thoracic aortas of *Usp20*^-/-^ and WT mice that were matched by age and sex. For a single line of primary ECs, aortas from 2-3 mice per genotype were isolated and cultured as reported previously (54). The immortalized MCEC line CLU510 was obtained from Cedarlane (Cedarlane/CELLutions Biosystems). The MCECs were grown in 150 mm cell culture plates containing Dulbecco’s Modified Eagle Medium (DMEM) supplemented with 10% fetal bovine serum (FBS) (v/v) and 1% (v/v) penicillin/streptomycin. 0.05% trypsin/EDTA (GE Life Sciences) was used to culture the cells.

### Adenoviral transduction

Recombinant adenoviruses (AdV) expressing HA-tagged USP20 WT (USP20-WT), USP20 dominant-negative (USP20-DN), USP20 phospho-mimetic (USP20-S334D), or USP20 phospho-resistant (USP20-S334A) mutants and GFP (control) were generated in our lab according to our previously published protocols (12,13,24). MCECs were cultured in media with low-serum (0.2%) for 6 h before infection with adenoviruses at a multiplicity of infection of 100 for 9 hours, after which they were serum-starved for 2 hours before conducting further analyses, namely, western blotting, spheroid-based sprouting, and scratch wound healing migration assays. To determine transduction efficiency, we processed parallel plates of adenovirus-infected cells for evaluating GFP fluorescence and HA epitope immunostaining, and conducted image analyses by confocal microscopy (23).

### Proinflammatory cytokine stimulation

MCECs that were transduced with recombinant adenovirus were stimulated with 10 ng/mL IL-1β for 15 minutes, after which the total protein was extracted with 2× SDS-PAGE protein sample buffer. The mouse aortic primary ECs were serum-starved overnight and subsequently stimulated with 10 ng/mL IL-1β for 5 to 60 minutes before protein extraction with 2× SDS-PAGE protein sample buffer.

### Western blotting

To determine the level of phosphorylation of the NF-κB subunit p65 on Serine 536 (phos-p65) in MCECs or mouse aortic primary ECs, immunoblots were performed using the total protein extracted in 2× SDS-PAGE protein sample buffer. Specific protein bands were separated by 10% SDS-PAGE gels and subsequently transferred to nitrocellulose membranes. The membranes were blocked with 5% (w/v) dried skim milk in TTBS buffer [0.2% (v/v) Tween 20, 10 mM Tris-Cl (pH 8.0 at 25°C), 150 mM NaCl]. The blocked membranes were incubated with phos-p65, β-actin or GAPDH antibodies overnight at 4°C. Membranes were washed three times with TTBS and incubated with the appropriate secondary antibodies for 1 hour at room temperature. Protein bands were detected with enhanced chemiluminescence (SuperSignal West Pico Reagent, Pierce) using a charge-coupled device camera system (Bio-Rad Chemidoc-XRS). Densitometry of protein bands was performed with Image Lab software (Bio-Rad).

### Immunofluorescence staining and confocal imaging

Immunofluorescence staining and confocal imaging of the cultured MCECs was performed according to our previously published protocol (12). MCECs transduced with recombinant adenoviruses were seeded on polyD-lysine-coated 35-mm glass-bottom plates. 48 hours post-transduction MCECs were fixed with 5% formaldehyde diluted in DPBS, and subsequently washed three times with DPBS. The fixed MCECs were permeabilized with 0.01% Triton X-100 in DPBS containing 2% BSA for 30 mins and incubated with the appropriate primary antibody overnight at 4°C. The next day, MCECs were washed with DPBS three times and incubated with the cognate secondary antibody as well as DAPI in a dark chamber for 1 hour at room temperature (RT). The stained MCECs were visualized with Zeiss LSM-510 META confocal microscope using a 63× oil immersion objective.

### Scratch wound healing migration assay

MCECs were grown to form a 90% confluent monolayer in 6-well plates containing DMEM supplemented with 10% FBS and 1% P/S. The MCEC monolayer was scratched with a 1 mL pipette tip to make “wounds,” and the medium was aspirated immediately to remove floating MCECs that had detached from the wounds made in the monolayer. The plates were washed once with DPBS, and 2 mL fresh DMEM supplemented with 10% FBS and 1% P/S was added in each well of the 6-well plates. If needed, inhibitors such as TPCA were added at this step. The images of the wounds were captured at different time points (0–16 h) by a Zeiss Axio Observer Z1 microscope with 2.5× magnification. At least 10 images per monolayer for each experiment were taken from each experimental group. Wound closure was quantified from wound width measurements using the ImageJ software. The extent of MCEC migration was calculated by the formula: 100 × (wound width at 0 h – wound width at indicated time point) / (wound width at 0 h) (55).

### Spheroid-based sprouting assay

We diluted 80,000 MCECs in 4 mL of DMEM supplemented with 10% FBS, 1% P/S, and 20% Methocel. The MCEC suspension was transferred to a sterile multichannel pipette reservoir and immediately 25 μL of MCEC suspension was pipetted as drops onto a 150 mm cell culture plate with the help of a multichannel pipette. There were approximately 400 cells in each 25 µL drop. The plate was incubated upside down in a CO_2_ incubator for 24 hours for spheroid formation. The hanging drops containing the spheroids were washed off with 10 mL PBS by pipetting them using a 10 mL serological pipette. The spheroid suspension was immediately transferred to a 15 mL conical tube and centrifuged at 200 ×*g* for 5 min. The supernatant was aspirated very carefully and 2 mL ice-cold Methocel supplemented with 20% FBS was added into the tube. Separately, a 4-mL collagen working solution (1 mg/mL) was prepared by adding the required volume of type I collagen in ice-cold DMEM supplemented with 10% FBS and 1% P/S. The pH of the collagen stock solution was adjusted between 7.0 and 7.4 by adding NaOH, and the stock solution was maintained on ice. The ice-cold Methocel containing the spheroids was mixed with the collagen stock solution, and immediately, 250 μL of the spheroid suspension was pipetted in each well of a 24-well cell culture plate. The plate was incubated in a CO_2_ incubator for 1 hour to polymerize and solidify the collagen. 250 µL of DMEM supplemented with 10% FBS, 1% P/S, and 30 ng/mL VEGF was added in each well of the 24-well plate containing the collagen matrix with spheroids embedded in it. If needed, inhibitors such as TPCA were added at this step. The plate was then incubated in a CO_2_ incubator for 48 hours. The images of sprout growth from the spheroids embedded in the collagen matrix were captured by a Zeiss Axio Observer Z1 microscope with 10× magnification. The number of EC sprout branch points (nodes) in a sprouted spheroid were quantified with NIH ImageJ.

### Aortic ring sprouting assay

The aortic ring sprouting assay was performed as described previously (56). Briefly, thoracic and abdominal segments of the aortas were isolated from age- and sex-matched WT, *Usp20*^-/-^, *Usp20*-S334A, and *Usp20*-S334D male and female mice. The isolated aortas were immediately washed twice with Dulbecco’s Phosphate Buffered Saline (DPBS) containing calcium and magnesium to remove residual blood, and then transferred to a 100-mm plate containing complete DMEM supplemented with 10% FBS and 1% P/S. Under a stereo microscope and with sterile technique, the residual fat from the aorta was removed with tweezers. Additionally, the side branches of the aorta were removed with a scalpel. The cleaned aorta was immediately transferred to a 100 mm plate containing complete DMEM supplemented with 10% FBS and 1% P/S. The cleaned aorta was dissected into small ring-like fragments with an approximate length of 0.8 mm. From a single aorta, 42–48 aortic rings were made. The aortic rings were transferred to a 6-well plate containing DMEM supplemented with 0.2% FBS and 1% P/S and incubated overnight in a CO_2_ incubator. The next day, a collagen stock solution was prepared in a 50 mL tube by adding 1 mg/mL collagen in ice-cold Opti-MEM supplemented with 10% FBS and 1% P/S. The pH of the collagen stock solution was adjusted between 7.0 and 7.4 by adding NaOH, and the stock solution was maintained on ice. 200 µL of the ice-cold collagen stock solution was pipetted into each well of a 24-well plate, and immediately, 7 aortic rings were planted in each well on top of the collagen matrix with the help of a sharp-tipped tweezers. The aortic rings were planted on the collagen matrix in such a way that the luminal axis of each aortic ring was perpendicular to the bottom of the well. The plate was then incubated in a CO_2_ incubator for 1 hour, and subsequently, 250 µL of Opti-MEM supplemented with 10% FBS, 1% P/S, and 30 ng/mL VEGF was added to each well of the 24-well plate containing the collagen matrix with aortic rings embedded in it. At this stage, the desired drug (for example, TPCA-1) was added to the Opti-MEM. The plate was then incubated in a CO_2_ incubator for 5 days, and then images were captured by a Zeiss Axio Imager Z2 Upright Microscope with 5×/0.16 Plan-Apochromat objective. The sprouting area of each aortic ring was captured by digital photography and subsequently quantified by an observer blinded to the genotype and treatments of each specimen. Sprouting area was quantified in ImageJ as the total area enclosed by the outer boundary of the sprouts minus the area of the central aortic ring and expressed as percentage sprouting area, as described previously (30,31,56).

### Proteome profiler analyses

The expression levels of angiogenesis-related proteins were determined with the Proteome Profiler Mouse Angiogenesis Array Kit (R&D Systems, Cat. No. ARY015) according to the manufacturer’s protocol. To extract proteins from the sprouted aortic rings, the wells of the 24-well plates containing the aortic rings were washed twice with PBS (with Ca^2+^ & Mg^2+^), and 150 µL RIPA buffer (12) was added in each well of the plates. The plates were incubated on a shaker overnight at 4°C. Supernatants were collected from the wells, and protein concentrations were measured with the BCA protein assay kit. Equivalent protein amounts of experimental samples were used for the angiogenesis proteome profiling assay.

### Statistical Analyses

All assays were repeated at least three independent times, and the data from all experiments were averaged and presented as means ± SD. Data were assessed for normality using the Shapiro–Wilk test. A t-test was performed to determine statistical significance between two groups. To compare more than 2 groups, we used one-way or two-way ANOVA followed by Holm-Šídák’s post hoc test for multiple comparisons. All statistical tests were performed using the GraphPad Prism 10 software (GraphPad, Inc). The level of statistical significance was set at *p* < 0.05.

## Acknowledgments

The authors thank Yushi Bai, Kamie Snow and Jaimie Dazio for their help in managing mouse lines used in this work.

## Author contributions

Conceptualization: B.R., N.J.F., and S.K.S. Investigation: B.R. and J.W. Formal Analysis: B.R., A.B., R.J., and S.K.S. Writing—original draft: B.R. and S.K.S. Writing— reviewing and editing: B.R., N.J.F., and S.K.S. Visualization: B.R., N.J.F., and S.K.S. Project Administration: S.K.S.

## Data and materials availability

All data needed to evaluate the conclusions in the paper are present in the paper or the Supplementary Materials.

## Supplementary Materials

This article includes Supplementary Figures S1–S3

