## Supplementary figures for "Ubiquitin-specific peptidase 20 constrains endothelial cell activation and angiogenic sprouting through NF-κB inhibition"

**Inflammation-induced endothelial cell activation and angiogenic sprouting are downmodulated by ubiquitin-specific peptidase 20**

Bipradas Roy[1], Jiao-hui Wu[1], Richard Jiang[1], Annie Bao[1], Neil J. Freedman[1, 2, 3] & Sudha K. Shenoy[1, 2, 4]

From the Departments of [1]Medicine (Cardiology) and [2]Cell Biology,
Duke University Medical Center, Durham, North Carolina 27710

### Supplementary Figures

**Figure S1**

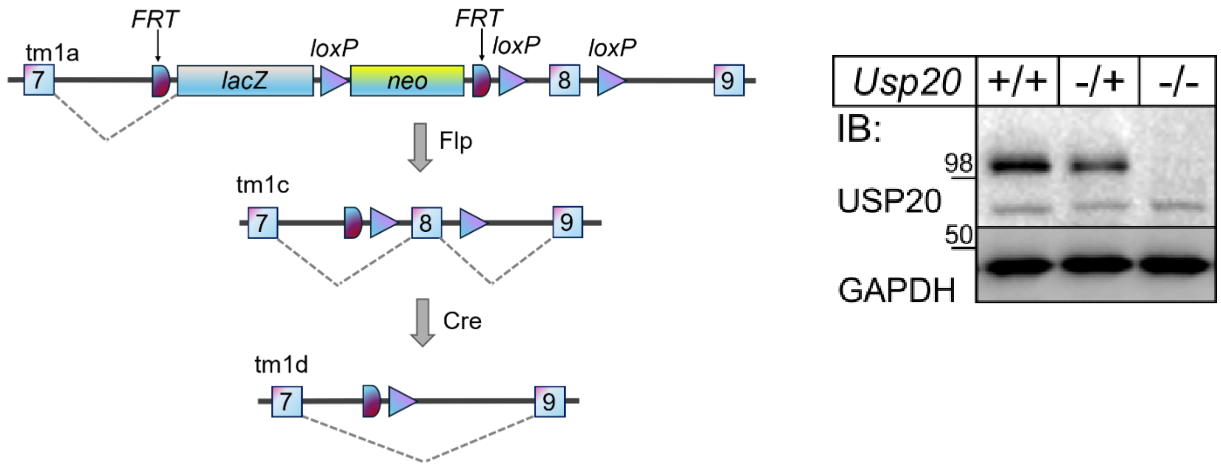


**Figure S1**. ***Usp20*$^{-/-}$ mouse generation.** Schematic shows the targeting vector design that contains knockout-first allele (tm1a) and the breeding steps that produce *Usp20*$^{-/-}$. The schematic is adapted from a model presented by KOMP and EUCOMM Breeding Strategies. Western blots show cardiac lysates of C57BL/6-congenic mice of the indicated genotype that were immunoblotted serially for USP20 and GAPDH (as a loading control). Shown are results from a single experiment, representative of 3 performed with distinct mice of each genotype.

**Figure S2**

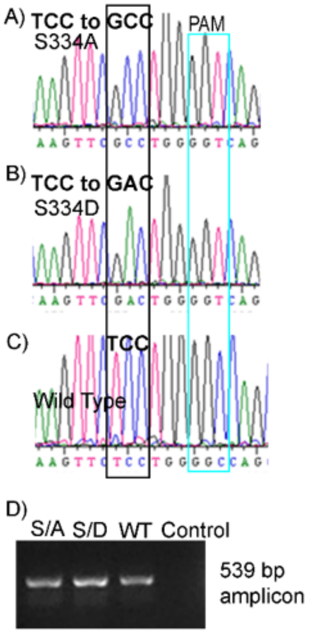


**Figure S2**. ***Usp20*-S334A and -S334D CRISPR/CAS9 gene-edited mice.** Panels A and B show sequence chromatograms of USP20 mutations: TCC to GCC, (Ser to Ala), TCC to GAC (Ser to Asp) confirmed by founder analysis and allelic subcloning. Panel C shows the WT sequence TCC (Ser334). Panel A and B also show the intended silent mutation (GGC to GGT) in the repair oligo to disrupt the Protospacer Adjacent Motif (PAM) sequence. (D) DNA gel showing the 539 bp amplicon of USP20 exon 9 region that contains the targeted site Ser334.

**Figure S3**

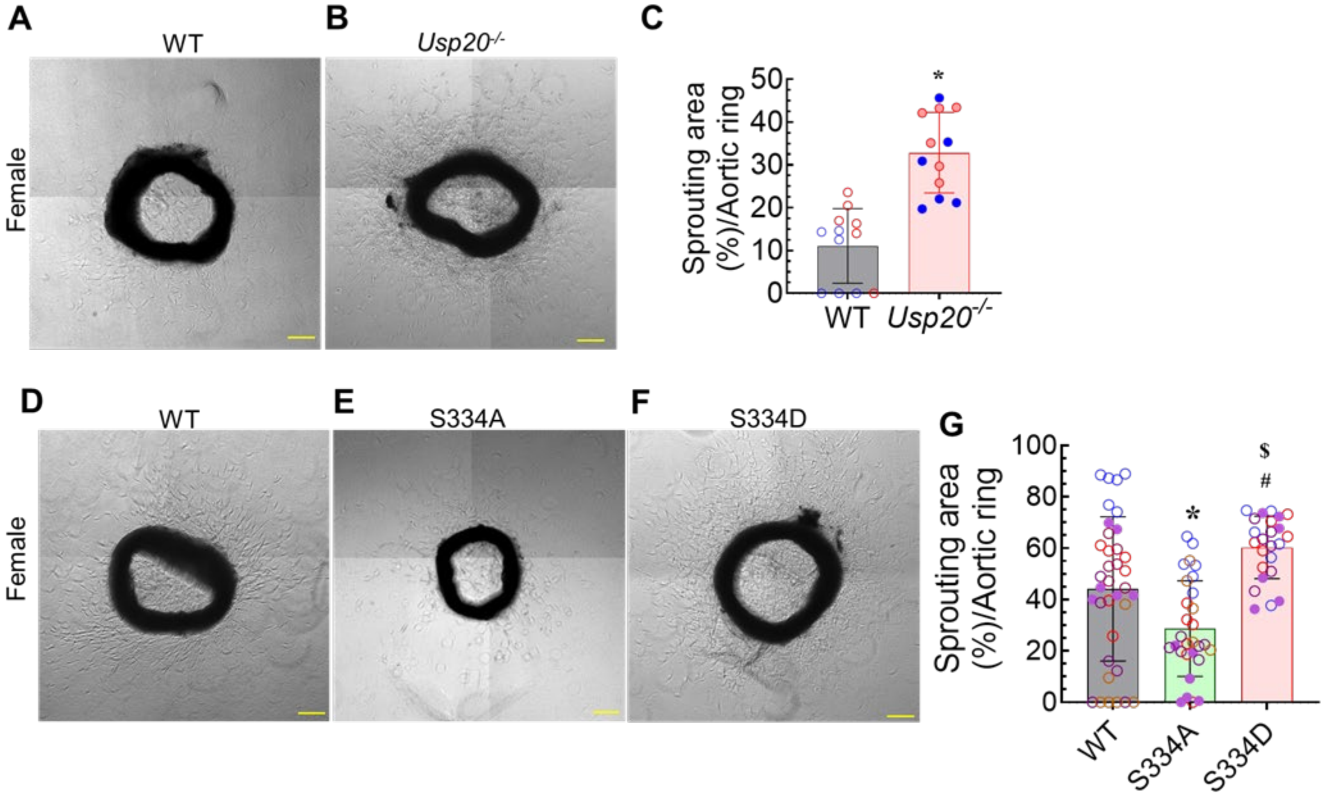


**Figure S3. USP20 activity constrains angiogenesis.** Aortic ring microvessel sprouting assays were performed with aortic rings from WT and *Usp20*$^{-/-}$ mice as in Fig 6, except that the mice were females. (A-B) Representative phase-contrast images of angiogenic sprouting are shown. Scale bar = 200 µm. (C) Angiogenic sprouting was quantified as in Fig 6. Data are presented as mean ± SD. Aortic rings were obtained from 2 WT and 2 *Usp20*$^{-/-}$ female mice. Compared with WT: *, $p < 0.01$ (t test). (D–F) Aortic ring microvessel sprouting was performed as in panels A and B, but with aortic rings from 3-month-old female WT, *Usp20*-S334A (phospho-resistant), and *Usp20*-S334D (phospho-mimetic) mice. Scale bars = 200 µm. (G) Angiogenic sprouting was quantified as in panel C. and is presented as mean ± SD. Aortic rings were obtained from 6 WT, 3 *Usp20*-S334A, and 3 *Usp20*-S334D female mice. Compared with WT: *, $p < 0.05$; compared with USP20-S334A: #, $p < 0.01$ (one-way ANOVA followed by Holm–Šídák's multiple comparisons test). In panels C and G each mouse represents 6 dots with the same color in each bar; each dot represents the average sprouting area of 4-7 aortic rings.
